# Long-term high throughput agitation culturing with real-time metabolic profiling

**DOI:** 10.64898/2025.12.28.696554

**Authors:** David Lenzen, Sarah Holbrook, Da-Han Kuan, Qinglan Ling, Cheng-Han Tsai

## Abstract

Cellular metabolism relies on the dynamic coordination between glycolytic flux in the cytosol and oxidative phosphorylation (OXPHOS) within the mitochondria. To study the metabolic profiles of cells, researchers apply a monitoring system for measurements of critical parameters, e.g., pH and dissolved oxygen (DO), to understand underlying energy production tendencies, dictating the performance, resilience and growth of cells. However, implementing sensitive, non-invasive sensors into long-term culturing environments remains a technical bottleneck. Here, we describe the DolphinQ bioanalyzer, a novel culturing platform designed for high-throughput, real-time monitoring of cellular metabolism states under physiologically relevant conditions. We validate the system across multiple cell types and experimental set-ups, demonstrating its ability to resolve subtle metabolic shifts that are typically obscured in end-point assays. Notably, we utilize the system to characterize the metabolic impact of heteroplasmy in a mitochondrial disease model with affected ATP synthase. Our results underscore the utility of continuous, minimally disruptive monitoring for revealing the complexities of cellular metabolic adaptation. The DolphinQ framework therefore offers a robust tool for optimizing culture conditions across a wide range of applications and advancing fundamental research into metabolic flux and mitochondrial dysfunction.

## 1. Introduction

Mitochondria have a double-membrane structure and circular genome which suggests a deep evolutionary history and multifaceted physiological roles [^1^]. This complexity contrasts with the long-forged concept that mitochondria are merely the ‘powerhouse of the cell’, which is rooted in their prominent function of aerobic respiration to generate the majority of ATP and essential metabolites needed for survival and growth [^2^]. Cellular energy production primarily involves two interconnected processes: cytoplasmic glycolysis and mitochondrial respiration. Glycolysis can occur in either aerobic or anaerobic conditions, yielding a net of 2ATP, 2NADH, and two molecules of pyruvate per glucose. Under an oxygen-deprived state (hypoxia or anoxia), pyruvate is converted to lactate in the cytoplasm to regenerate NAD^+^, thereby allowing glycolysis to continue its limited production of ATP [^3^]. Whereas with sufficient oxygen (normoxia), this pyruvate enters the mitochondria, where it is first converted to acetyl-CoA to fuel the subsequent Krebs cycle (or citric acid cycle). This cycle yields the electron carriers NADH and FADH^+^, which then donate their electrons to the Electron Transport Chain (ETC). At the terminal step of oxidative phosphorylation (OXPHOS), the established proton gradient drives the generation of the majority of total ATP production [^4^]. The capacity of a cell to modulate these pathways in response to environmental or genetic stress, termed “metabolic flexibility”, is a fundamental determinant of health and disease [^5, 6, 7^].

Mitochondrial diseases, which rank among the most frequent inborn errors of metabolism, generally stem from OXPHOS dysfunction [^8, 9, 10^]. This impairment is driven by mutations in mitochondrial (mtDNA) or nuclear (nDNA) DNA that disrupt the central pathway for ATP synthesis. The resultant energy deficiency is more likely to affect high-energy-demanding organs, such as the brain, liver and muscles [^11, 12^]. This OXPHOS insufficiency negatively affects cellular viability not only through insufficient energy supply, but may also disrupt calcium signaling and normal reactive oxygen species (ROS) levels [^13, 14^]. These combined molecular effects can induce cell death via apoptosis or necroptosis, causing organ damage and a broad spectrum of systemic symptoms, including fatigue, exercise intolerance and neurological or multi-systemic disorders. Consequently, the specific clinical presentation and disease severity vary widely based on the extent of the mitochondrial impairment [^9, 15^]. OXPHOS dysfunction and other mitochondrial stress trigger the integrated stress response (mt-ISR) to orchestrate comprehensive metabolic rewiring supporting cellular survival [^16, 17^]. These extensive studies in cell and animal models show that the mt-ISR represents a fundamental cellular adaptation mechanism with dual protective and pathological roles. Investigating how cells respond to mitochondrial dysfunction is key to understanding the physiological function of the organelle and ultimately clarifying the pathogenesis of mitochondrial diseases. Thus, a device or utility platform that allows for the dynamic monitoring of both OXPHOS and glycolysis would help achieving this goal, and potentiate therapeutic and diagnostic strategies for mitochondrial diseases, as well as other prevalent medical conditions like cancer, diabetes, and autoimmune diseases [^10, 18^].

The Agilent Seahorse Extracellular Flux (XF) Analyzer, which utilizes high-sensitivity, high-throughput, fluorescent optical-chemical sensor technology, remains the paradigmatic platform for OXPHOS and glycolytic activity [^19^]. The bioanalyzer directly measures dynamic changes in both dissolved oxygen concentration and proton flux within the cell culture supernatant. These readouts are subsequently converted into the oxygen consumption rate (OCR) and the extracellular acidification rate (ECAR), providing an assessment of ATP production capacity to evaluate the relative balance between mitochondrial respiration and glycolysis [^20^]. In a typical inhibition assay, preformulated compounds like oligomycin, FCCP, and the rotenone/antimycin A are automatically injected into the microtiter wells to reveal aspects of the target cells’ bioenergetic trajectory [^21^]. Despite numerous upgrades and revisions, the platform retains limitations that continue to restrain research efforts. Most notably, the Seahorse assay duration is typically limited to 1-2 hours and is based on fixed time-point data collection which cannot fulfill continuous, long-term observation. For example, a comprehensive survey of cellular responses to a metabolic modulator often requires uninterrupted monitoring spanning from overnight to several days. This is most needed for cell types or biological models requiring prolonged growth durations to achieve maturity, such as neuron dendritic outgrowth studies, or complex organoid systems. During growth, cells undergo dynamic metabolic reprogramming that may occur on rapid or gradual timescales, making such changes difficult to resolve using endpoint-based assays[^22, 23^].

An alternative approach to the Seahorse flux analyzer lies in microfluidic organ-on-a-chip technology. As an example, Bavil et al. developed one such microfluidic platform, which integrated oxygen-sensitive phosphorescent microprobes to achieve real-time oxygen uptake monitoring during the 28-day culture of HepG2/C3A 3D cell aggregates under stable oxygen supply and flow stimulation [^24^]. The time-course profile unveiled metabolic changes of the cells following different drug treatments. Notably, the embedded electrochemical sensor allowed the researchers to identify and quantify these metabolic changes, attributing contributions to OXPHOS, glycolysis, and glutaminolysis. While equipped with these impressive improvements, the system is nonetheless constrained by its unsatisfactory throughput. Furthermore, manipulating microfluidic chips remains labor-intensive and usually requires educated personnel, collectively hindering broader adoption.

To address the technological limitations of the two approaches above, we developed the DolphinQ bioanalyzer. A novel cell culturing platform that enables continuous, long-term monitoring of live cells under flexible conditions. The system accommodates four independently controllable chambers compatible with standard 24- and 96-well culture plates. During culture, steady pneumatic mixing improves nutrient and oxygen distribution while simultaneously reducing the shear stress often seen in other mixing types, thereby maintaining cell viability over extended periods [^25^]. This mixing strategy ensures that DO and pH detection via sensors at the well bottom more accurately reflect the true metabolic nature of the cells than is possible under static or irregular culture conditions. With the cells being able to grow in the media and environment they are accustomed to, this format eliminates one of the key physiological uncertainties of other metabolism assays, which keeps cells in often unbuffered environments, potentially affecting their health and morphology[^23^]. Together, DolphinQ offers not only a comprehensive experimental narrative but also high-resolution analysis toward the dynamics and strategies of cellular adaptation to mitochondrial performance.

In this study, we show the architecture of the DolphinQ bioanalyzer and validate the system with three disease-relevant cell lines. We first detailed the determination of oxygen transfer coefficient (kLa) using the static gassing out method. The kLa value is prerequisite for accurately translating DO into the OCR under stable mixing and environmental conditions. We then illustrated the mathematical and computational frameworks employed to convert these DO and pH measurements into the respective metabolic metrics, oxygen consumption rate (OCR) and extracellular acidification rate (ECAR). We validated baseline metabolism profiling performance using the commonly used human adenocarcinoma cell line A549. Groups consisted of different seeding densities of these highly metabolically active cancer cells. Subsequently, we present data from a collaboration with Qinglan Ling’s Lab (Horae Gene Therapy Center, University of Massachusetts Chan Medical School) investigating a Leigh Syndrome cellular disease model. The DolphinQ system was utilized to monitor A375 human melanoma cells [^26^] and human primary fibroblasts bearing various degrees of mtDNA mutations. Our data demonstrated that DolphinQ can distinguish distinct cellular metabolic patterns of mitochondrial dysfunction, thereby highlighting its potential as a valuable platform for research inquiring cellular mitochondrial function and metabolic state changes.

## 2. Materials and Methods

### 2.1. Determination of Volumetric Oxygen Transfer Coefficient (kLa)

The volumetric oxygen transfer coefficient (kLa) was quantified using the static gassing-out method using the DolphinQ system (Leadgene Biosolutions (Former Cytena Bioprocess Solutions), Taiwan). A 24-well plate with integrated dissolved oxygen (DO) and pH sensors, together with the corresponding 24-well DolphinQ lid (Leadgene Biosolutions, Taiwan), was used, as well as a 96-well plate with integrated DO sensors and the corresponding 96-well DolphinQ lid (Leadgene Biosolutions, Taiwan). Dulbecco’s Phosphate-Buffered Saline (DPBS; Corning, USA) was used as the working solution, with 1400 μL added to each well in all kLa measurements. Measurements were programmed in the engineering software to record signals at one-minute intervals, at mixing rates set to 2, 5, 10, 25, or 50 seconds per cycle, and the chamber temperature maintained at 37 °C.

### 2.2. Cell Lines

The base A549 cell line used for seeding density experiment was purchased from the Taiwanese Bioresource Collection and Research Center (BCRC). A549 cells were cultured in high glucose Dulbecco’s modified Eagle’s medium (Gibco Cat#11960051), 10% fetal bovine serum (Gibco Cat#A3160601), 1x sodium pyruvate (Corning Cat#25000CI), 1x L-glutamine (Corning Cat#25005CI), 1x Pen-Strep (Corning Cat#30002CI). The A375 cells were obtained from Prashant Mishra’s lab at UT southwestern Medical Center [^26^]. Human Fibroblasts cell lines were purchased from the Coriell Institute (GM13411). Cells were cultured and maintained in phenol red-free DMEM (Sigma Cat#D5030) supplemented with 10% fetal bovine serum, 4.5 g/L D-(^+^)-Glucose (Sigma Cat#G7021), 3.75 g/L NaHCO_3_ (Gibco Cat#25080094), 1xGlutaMAX (Gibco Cat#35050061), and 1x sodium pyruvate (Gibco Cat#11360070), 1xNEAA (Gibco Cat#1140050), 1x Pen-Strep (Gibco Cat#15140122), and 0.05 g/L Uridine. Rho0 A375 cells and A375 cybrid cells with MT-ATP6 mutation were generated as described before and subsequently validated via quantitative PCR [^26^]. All culturing was done at 37 °C in a 5% CO^+^ humidified atmosphere.

### 2.3. MT-ATP6 heteroplasmy level determination in cultured cells

The heteroplasmy levels were evaluated using ddPCR and confirmed with Nanopore sequencing using services from Plasmidsaurus, Inc (Louisville, USA). Genomic DNA was isolated from cells using Monarch Spin gDNA Extraction Kits (New England Biolabs Cat#3010L). The MT-ATP6 gene fragment spanning the mutation m.8993 site was amplified with the use of Phusion High-fidelity DNA Polymerase (New England Biolabs Cat#M0530L) with Forward primer-GCTTCATTCATTGCCCCCAC and Reverse primer-AGGCGACAGCGATTTCTAGG. PCR fragment was then cleaned with Monarch Spin PCR & DNA Cleanup Kit (New England Biolabs Cat#T1130L). Purified PCR products were sequenced with Plasmidsaurus through the Purified PCR sequencing service (Sup. Figure 2). CRISPResso2 was used to analyze the percentage of T to G mutation.

### 2.4. DolphinQ culture

Prior to the experiment, cells were seeded onto the DolphinQ sensor plate and cultured until sufficiently adherent. These cell types had sufficient natural adherence to not require additional coating for the mild mixing environment. However, multiple common coating strategies like Matrigel, Laminin, or Poly-L-Lysine have been evaluated and confirmed to be compatible with the sensors. Images confirming accurate seeding and baseline confluency were captured using the CloneSelect Imager (Molecular Devices, USA). Subsequently, the DolphinQ pneumatic mixing lid was placed on the sensor plate and that assembly then positioned atop the plate holder tray. 40 mL of ddH^+^O were added to the water tray surrounding the plate. A pre-warm experiment was run for 45-60 minutes, assuring identical culturing conditions across the entire plate before initiating the experiment run stretching between 3 and 5 days, depending on the experiment. Upon experiment completion, the sensing plate and lid were removed and taken for subsequent testing. The water tray was emptied, and chamber sterilized by mildly swiping off the residual condensation with a cotton swap, followed by a mild 70% ethanol swipe. Afterwards, the chamber was further sterilized with the integrated UV light function.

## 3. Results

### 3.1. System Design

Four independently controlled chambers (Figure 1a) accommodate standard 96- or 24-well plates and utilize the unique DolphinQ lid (Figure 1b) to introduce vertical mixing during the culture period. This specially designed lid is the key component of the chamber, securely docking to the well plate connectors (Figure 1c). If mixing isn’t required, each chamber can also be used as a standard static incubator, using a standard culture plate and lid. The mixing mechanism relies on an integrated switching valve to control the direction of gas flow, moving in or out of the chamber lid. When the valve directs gas inflow (pressure), the air is forced onto the liquid surface, expelling the media downwards (Figure 1d). Conversely, when the valve directs gas outflow (vacuum), air is drawn away, causing the liquid to be pulled upwards (Figure 1e). By delicately alternating between expulsion and suction, the system creates a continuous, reciprocating flow that effectively mixes the media. This pneumatic action simulates manual pipetting to homogenously distribute nutrients, promote effective aeration and minimize damaging shear stress. Gas composition can be adjusted via the application software. For example, the system can be set to supply 5% CO^+^ for standard cell culture or deliver atmospheric air/pure nitrogen for kLa determination, providing essential flexibility for different assay requirements.

**Figure 1.**
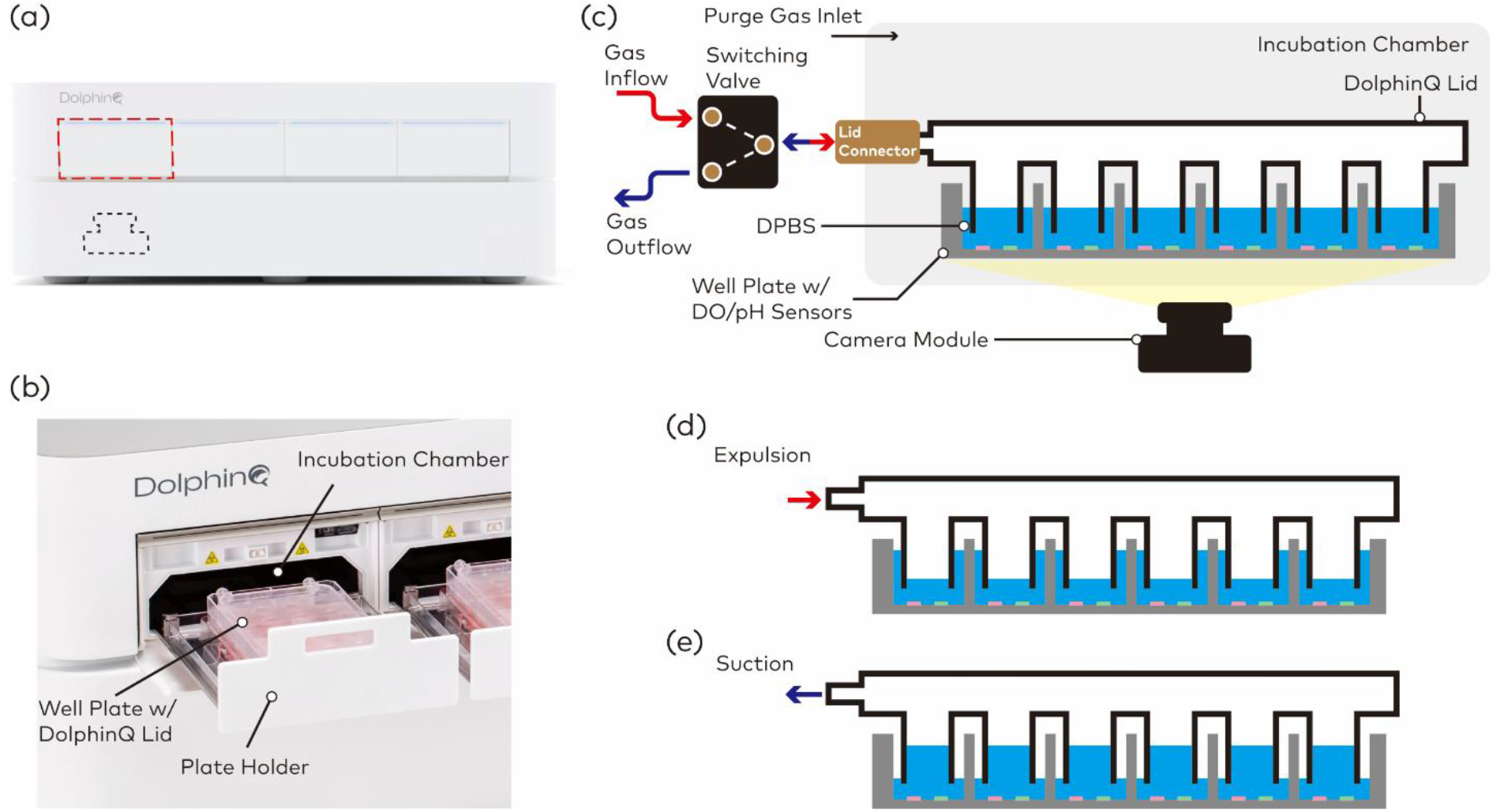
Images of (a) the DolphinQ system and (b) the interior components of each incubation chamber. (c) The schematic of the incubation chamber of DolphinQ. (d-e) The illustration showing the principle of introducing reciprocating mixing in the well plate.

Additionally, fluorescence-based DO and/or pH sensors are equipped for real-time monitoring in each well. A camera module is located beneath the chamber, capturing images of the entire plate every 10 minutes and then converting the fluorescence signal to quantitative DO and pH values. The DolphinQ software will collect and display DO and pH profiles on the dashboard. Due to the integration of mixing and sensing, DolphinQ can offer a non-destructive and non-invasive metabolism characterization during cell culture, particularly suitable for those cell models with slow metabolic transition.

### 3.2. Determination of Volumetric Oxygen Transfer Coefficient (kLa)

To convert the DO profile to oxygen consumption rate, the volumetric oxygen transfer coefficient (kLa) of the DolphinQ was first quantified. For this purpose, the static gassing out method was selected for kLa measurement [^27^]. As illustrated in Figure 2a, the method can be divided into three phases: calibration, oxygen deprivation, and oxygen dissolving. Prior to the measurement, the chamber temperature was set to 37 °C, and the mixing rate was fixed at 2, 5, 10, 25, or 50 seconds per cycle. In the calibration phase, the atmospheric air was supplied as both the driving and purge gas for sensor initialization. Once the sensors reached equilibrium state, the corresponding fluorescence signal was calibrated as 100% DO. During the oxygen deprivation phase, both gas flows were replaced with pure nitrogen to establish an oxygen-free environment. This causes a reduction of the dissolved oxygen concentration in the solution until the level was near zero. In the final oxygen-dissolving phase, the atmospheric air was reintroduced to allow oxygen to re-dissolve into the medium and increase the DO level over time. Importantly, the usually 10-minute imaging interval was shortened to 1-minute to achieve maximal sensitivity, and a more concisely observable oxygen-dissolving phase.

**Figure 2.**
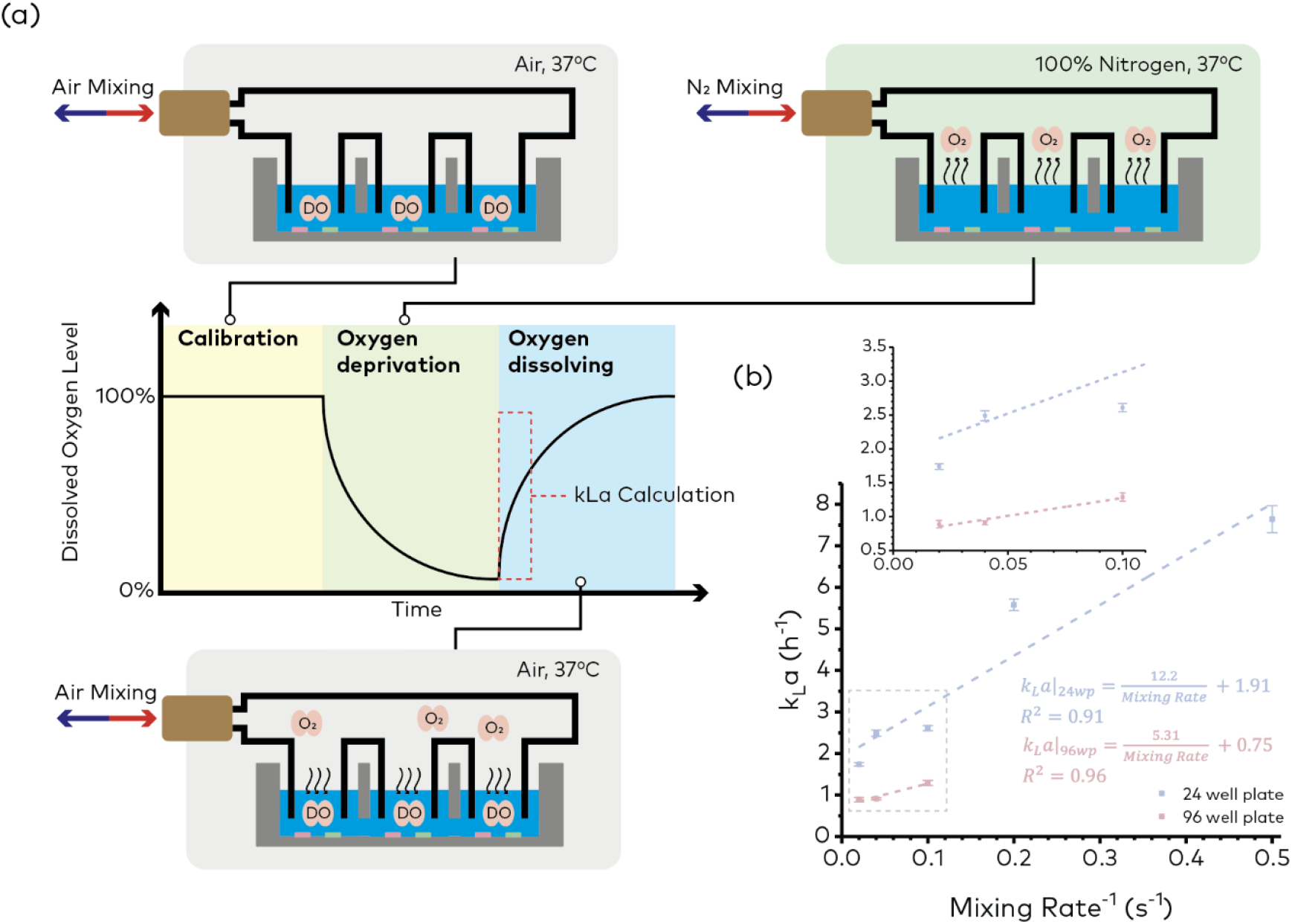
(a) The schematics showing chamber environment conditions and the expected dissolved oxygen profile of the static gassing out method for kLa determination. (b) The transfer functions between kLa and mixing rate under 24- and 96-well plate formats.

By analyzing the time-course dissolved oxygen profile during the oxygen-dissolving phase (highlighted with red dashed line in Figure 2a), the oxygen transfer coefficient of the corresponding mixing rate was obtained from calculating the slope of the linear regression of following equation: ln 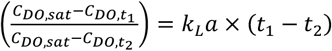, where *C*_*DO,sat*_ is the saturation DO concentration of the medium, *C*_*DO,t1*_ and *C*_*DO,t2*_ are the local DO concentration of the medium at *t*_1_ and *t*_2_. Figure 2b plots the calculated coefficients at different mixing rates and well plate formats. For the implementation of kLa conversion from mixing rate in the DolphinQ software, the line regressions between oxygen transfer coefficients and mixing rate in the 24- and 96-well plates were calculated as well.

### 3.3. Oxygen Consumption Rate (OCR) and Extracellular Acidification Rate (ECAR) Conversion

To minimize the conversion error resulting from the noise, the raw dissolved oxygen and pH profiles were first processed utilizing baseline correction followed by locally weighted scatterplot smoothing (LOWESS), shown in Figure 3. For each experiment, at least two wells were filled with pure medium as blanks on every plate. These blanks reflected the background signal resulting from the mild environmental fluctuation (e.g., temperature and CO2 concentration) and potential fluorescence signal drift. For DO, the baseline correction was performed by normalizing the blank to 100% DO level, while for pH, the blank was shifted to its initial value, and the same offset was applied to the corresponding sample signal. Subsequently, LOWESS was applied to reduce high-frequency random fluctuation, such as thermal noise. The algorithm performs locally weighted polynomial regression within a specific span to generate the best-fitting points. Data processing was implemented in the DolphinQ software using the ‘stats-lowess’ function from the JavaScript stdlib library. To preserve raw information while suppressing noise, the smoothing span f was set to 0.2, the smallest recommended value described in Cleveland’s work [^28^]. The smoothened profiles were then converted to oxygen transfer rate (OCR) and extracellular acidification rate (ECAR) based on the corresponding equations for DO and pH. The constant kLa for OCR conversion was obtained from the previous characterized kLa mixing transfer function.

**Figure 3.**
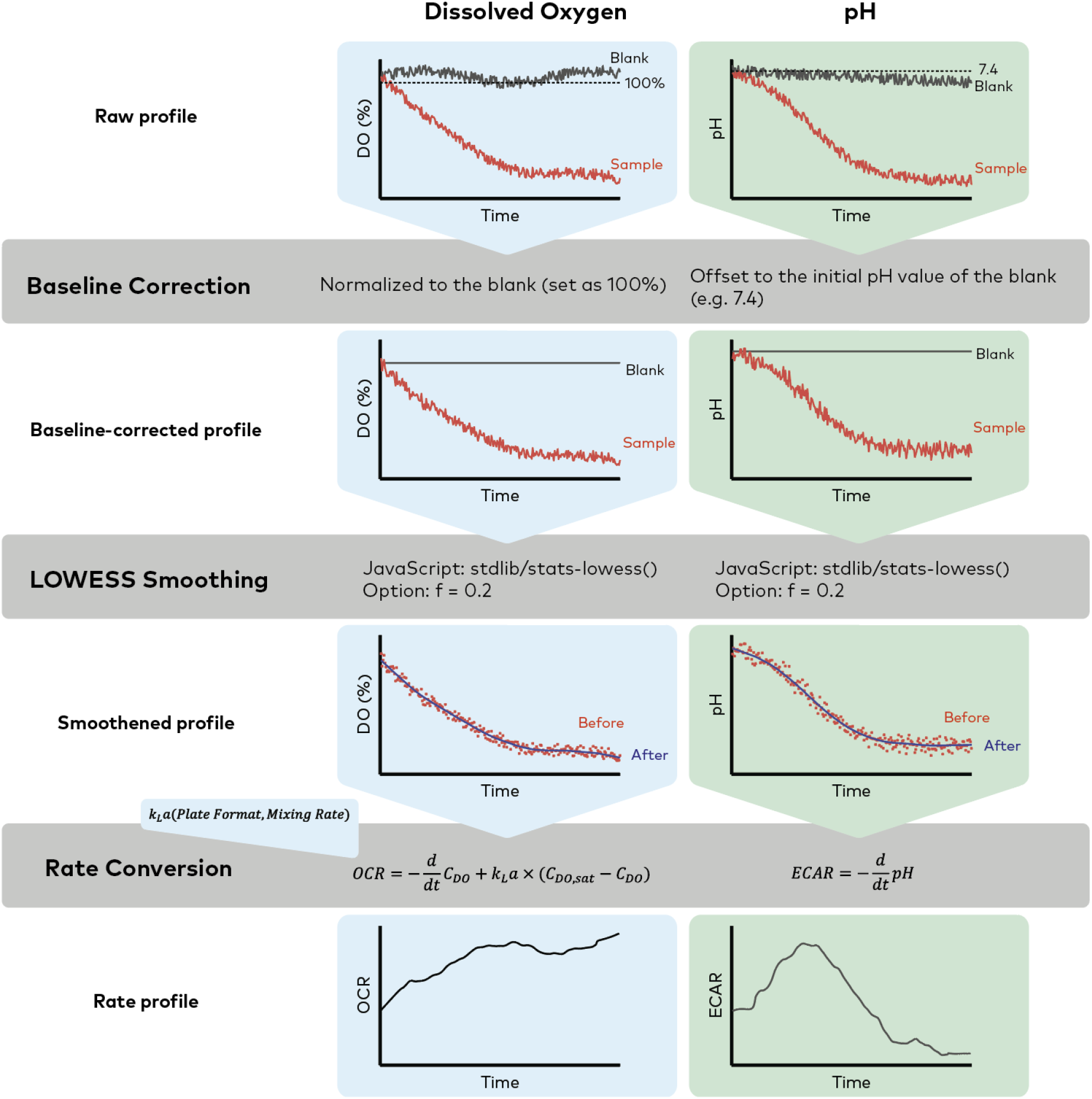
Steps for converting dissolved oxygen level (DO) to oxygen consumption rate (OCR) and pH to extracellular acidification rate (ECAR), including baseline correction, LOWESS smoothing, and rate conversion. The oxygen transfer coefficient (kLa) for OCR conversion is based on the result from kLa determination.

### 3.4 Assesment of DolphinQ baseline performance

The most prominent factors regulating metabolic paces of a cell population inside a microwell are initial seeding density, nutrient supply of the media, and the overall environments parameters such as temperature, mixing speeds and balances between dissolved gases, including O2 and CO2 [^29, 30^]. To assess the system’s consistency and accuracy, we designed a simple 4-group, 3-day seeding density experiment using adherent A549 human adenocarcinoma cells. As cells are often characterized by doubling time, the groups were chosen as a geometric progression from 5×10^4^ to 4×10^5^ cells/well. To confirm equal seeding density within each group, contrast imaging was done (Sup. Figure 1b-c). Four wells (A1, B4, C3, D6) were used as media blanks to assess baseline readout consistency across the plate (Sup. Figure 1a). These blanks were used as baseline correction standards as they represent both inner and outer well parameters. All groups were clearly separatable from one another and showed distinct glycolytic and aerobic metabolism strengths (Figure 4). Lower seeded groups took about one day to achieve comparable DO and pH readouts to their duplicate counterpart, which is explainable with A549’s doubling time of around 22 hours. Notably, with increased seeding density, the intra-group variance increased. This was to be expected, as even minor liquid handling variability could result in significant loss or addition of cell number at these highest seeding densities. Figures 4e-i show the run once more, however with the data baseline corrected to the media blanks and are shown in a weighted average format through an integrated software setting. Baseline correction can help remove pause or run start related spiking as the chambers become stabilized at the pre-selected culturing parameters. A key feature of the post processing algorithm is visible here; We can observe the effects of the LOWESS smoothening, in particular the early ECAR readouts where the lack of data points and therefore low confidence produces a generally flat readout. These tools allow for the full or partial removal of natural fluctuations and noise, such as seen within the first hour of the experiment or if the chamber had to be opened for treatment addition or media changes. This experiment aids as proof-of-concept towards future cultures and shows the benefits of baseline correction and adequate blank distribution.

**Figure 4.**
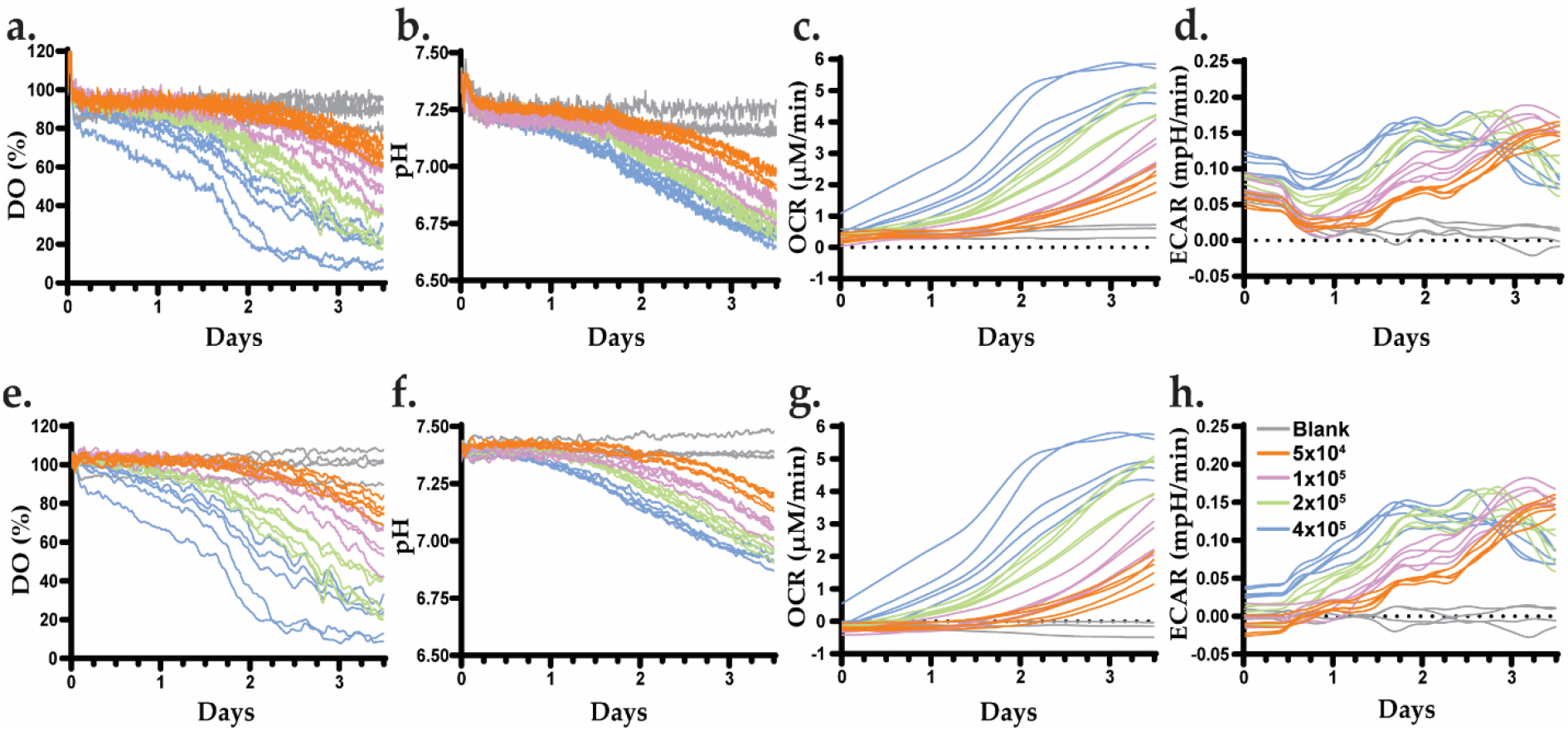
Proof-of-concept experiment for the use of the DolphinQ system in metabolic research. (a-d), A549 cells were used to create a cell seeding dependent metabolic profile. Left to right: Dissolved Oxygen percentage, Oxygen Consumption Rate (OCR) in μM/min, relative pH of media, and the Extracellular Acidification Rate (ECAR) in mpH/min are shown. n=5. (e-h), represents the identical run to a-d with weighted average data, baseline corrected to the blank wells.

### 3.5 Characerization of mitochondria disfuction in 24 and 96 well formats

One of the applications of the DolphinQ system is to profile cellular models of mitochondrial diseases. Mitochondrial diseases are long-term, severe, multi-systemic inherited disorders that affect one in every 5,000 individuals. Healthy mitochondrion requires co-operative effort of more than one thousand proteins, of which 99% are encoded by nuclear DNA (nDNA), while 1% is transcribed and translated inside mitochondria from mtDNA [Zech et al.]. Therefore, in a joined effort with Qinglan Ling’s Lab at the University of Massachusetts Chan Medical School, we demonstrated the capability of DolphinQ in characterizing metabolic activities represented by cell groups associated with disorder genotypes. First, a run containing 3 key groups of A375 melanoma cells (from here on called Wildtype (WT), Mutant (MUT, >80% VAF), mtDNA depleted (Rho0)) was performed in the 24-well sensing format (Figure 5a). Rho0 cells are strong experimental controls for metabolic cell culturing as they nearly exclusively rely on glycolysis to generate ATP due to their lack of mitochondrial DNA. This was consistently observable throughout the readout, which for simplicity is shown via group averages with Standard Error of the Mean (SEM). Rho0 cells presented near zero oxygen consumption (Figure 5b-c) and showed lower viability and growth after 3 days (not shown). Interestingly, although the untreated WT control cells exhibited the highest OCR, the mutant cell group displayed substantially higher ECAR measurements than both the Rho0 and WT groups. Those results support the prior hypothesis that cells with elevated levels of MT-ATP6 heteroplasmy largely upregulate glycolysis to maintain ATP sufficiency and promote survival. This metabolic shift, often compared to the Warburg effect of cancer cells, is commonly observed in cell models testing for metabolic flexibility. These findings were then reproduced in an identical experimental setting using 96-well plates (Figure 5f–h). The 96-well groups, having been seeded with a much smaller cell quantity (3×10^4^ vs 1×10^5^ cells/well), showed very comparable trends and had lower overall variance leading to a highly accurate and consistent readout. Interestingly, they yielded a more constrained total readout compared to the 24-well format. Although the qualitative result remained consistent, the visual representation and raw numerical values varied noticeably between the two plate formats. This scaling difference must be considered when designing experiments based on desired timeline and throughput.

**Figure 5.**
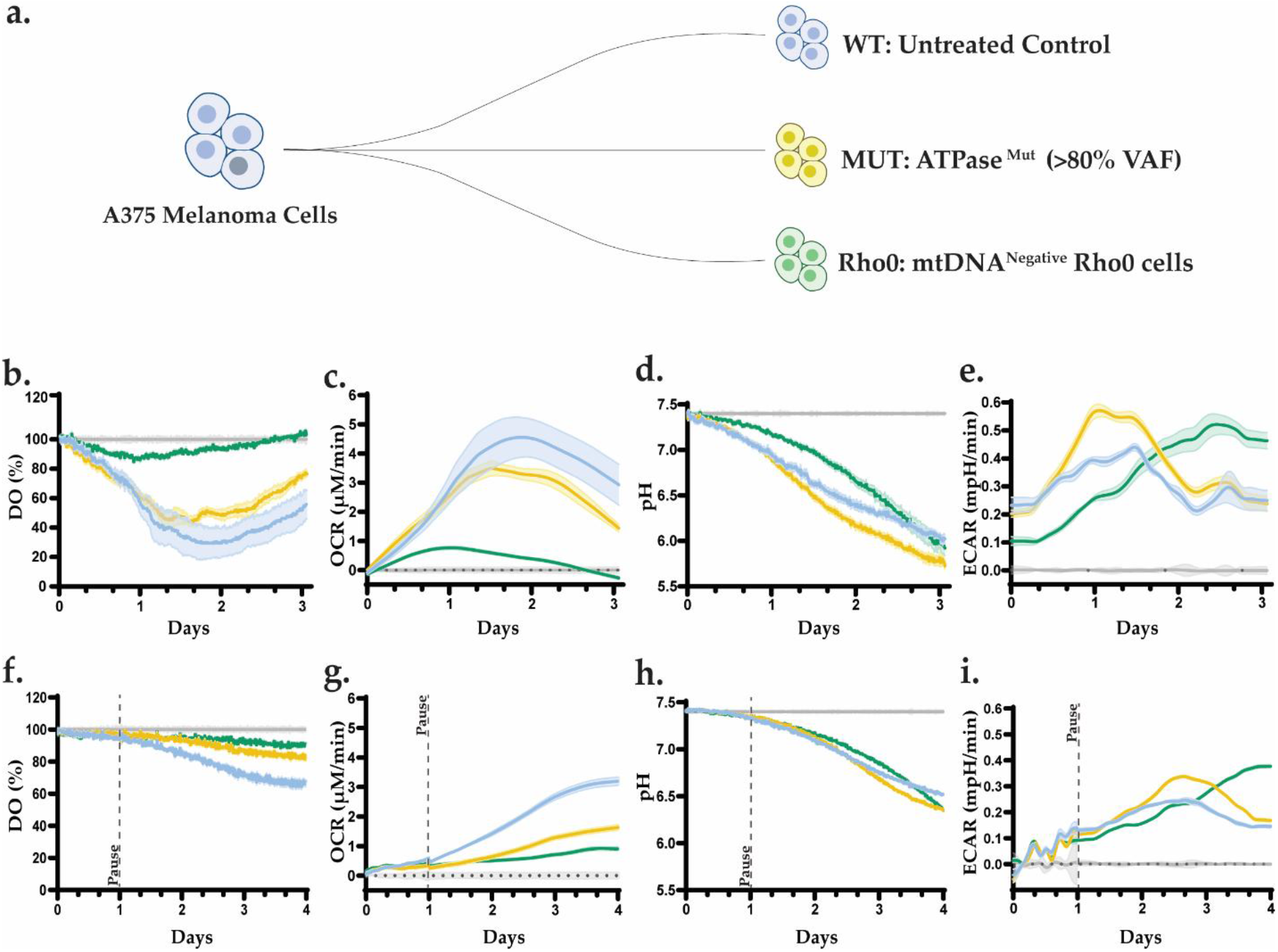
Mitochondrial genotype influences oxygen consumption, pH, and metabolic fluxes in A375 melanoma co-cultures. (a) A375 melanoma cells were used to create three groups of cell populations: Healthy Control (blue), ATPase LOF mutation harboring >80% mtDNA variant allele frequency (yellow), and mtDNA-negative Rho0 cells (gray). (b-e) Show the metabolic performance in the DolphinQ 24-well sensor plates. In order: Dissolved Oxygen percentage, Oxygen Consumption Rate (OCR) in μM/min, relative pH of media, and the Extracellular Acidification Rate (ECAR) in mpH/min are shown. All wells’ seeding density was 1×10^5^, n=5. (f-i) Show the metabolic performance in the DolphinQ 96-well sensor plates using the identical set up as seen in the previous 24-well experiment (n=7-8). All 96-well microwells seeding density was 3×10^4^, n=7-8. All data is baseline corrected to the blanks and represent mean ± SEM. Pauses indicated by dotted vertical lines.

### 3.6 Confirmation of metabolic disease profiles in human dermal fibroblasts

To further evaluate the DolphinQ capability in characterizing cellular metabolic flux changes under mitochondrial dysfunction, we used primary human fibroblasts, a cell type less metabolically active than the A375 melanoma cell line. Given the limited availability of cells harboring the mtDNA mutation, a 96-well format was chosen. Both Wild-Type (WT) and Mutant (MT) cells with m.8993T>G mutation were treated with the ATP synthase inhibitor oligomycin after one day of culture. Oligomycin treatment induced an immediate decline in OXPHOS and a concomitant spike in ECAR. (Figure 6). Notably, the WT cell group exhibited a greater shift toward glycolysis compared to the near homoplasmic MT. The markably lower total metabolic performance of these fibroblasts compared to A375, aids as a valuable example of how the system can maintain high accuracy even at low total dissolved oxygen changes.

**Figure 6.**
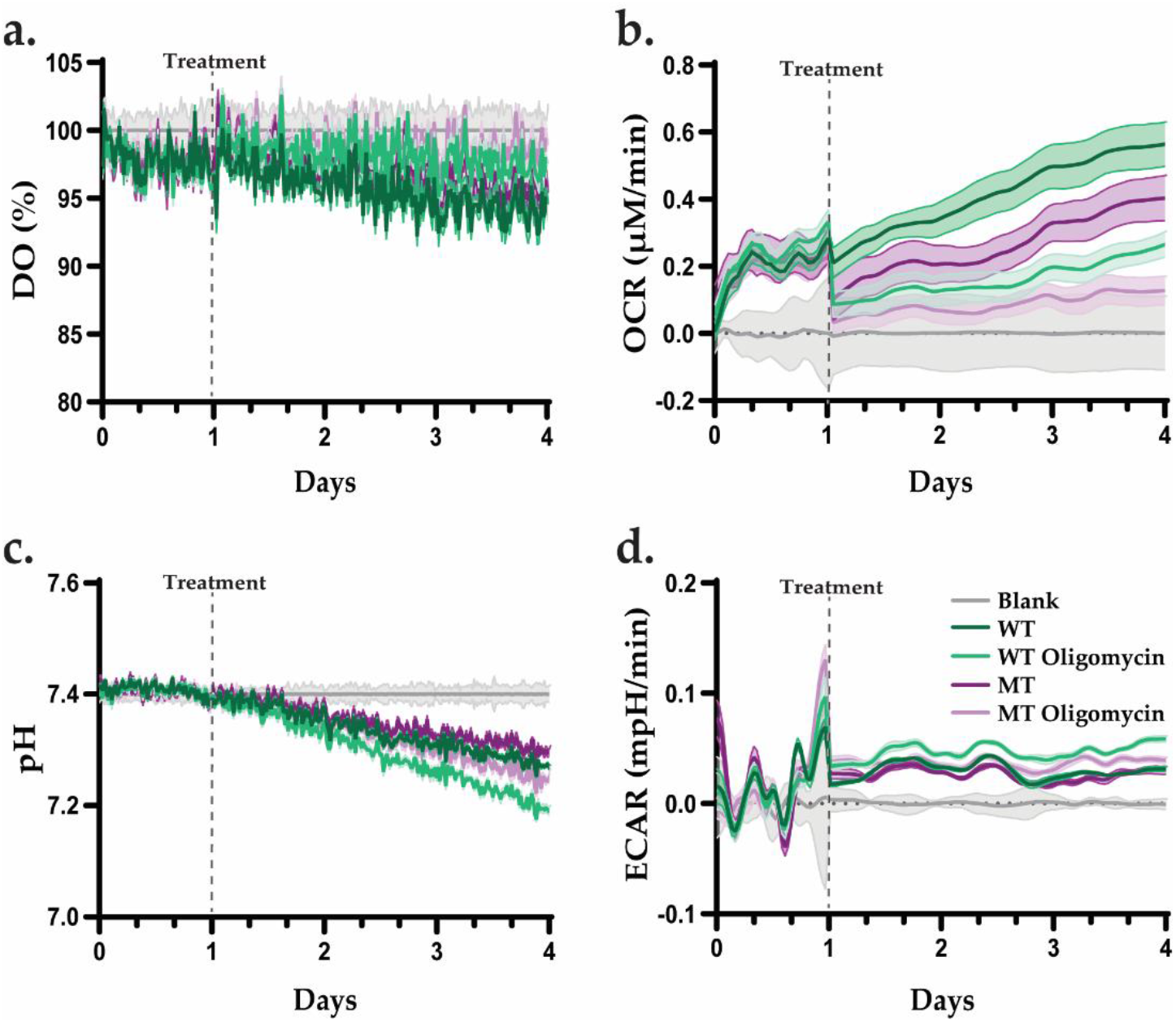
Show the metabolic performance of human dermal fibroblasts in the DolphinQ 96-well sensor plates when treated with the ATPase inhibitor Oligomycin. All wells’ seeding density was 3×10^4^, n=7-8. All data is baseline corrected to the blanks and represent mean ± SEM. Pauses indicated by dotted vertical lines.

## 4. Discussion

This study shows the technology and real-life application of a novel cell culture system capable of measuring the well-specific DO and pH of cultured cells in a long-term and non-invasive manner. While the positive effects of pneumatic mixing on cell health and OTR have been reported before, combining non-invasive DO/pH measurements with it allows for long term healthy growth and observation of multiple cell types [^25^]. The study was designed to focus on the mathematical and computational models supporting the device, which allow for consistent high-quality metabolic readouts spanning hours to weeks. The DolphinQ system provides researchers with 4 fully independent chambers capable of adjusting temperature, CO2, mixing speed and rhythm. To achieve consistent mixing, the system uses a special pneumatic airflow lid compatible with most commercial 24- and 96-well plates. During mixing, expulsion and suction cycles control the mixing speed and by doing so, affect the OTR of the culture (Figure 1). The OTR is vital to calculating the OCR of a solution and requires continuous homogenization of the fluid [^30, 31^]. To obtain an accurate OTR based on mixing speed, we performed volumetric oxygen transfer coefficient measurements using the static gassing out method (Figure 2). This method was chosen due to its well-established accuracy and ease of installation with the DolphinQ device [^27^]. The kLa was established using a linear regression model and calculated for each commonly used mixing strength (Table 1). The program, having been fed with adequate kLa values, is subsequently capable of interpreting the fluorescent DO signal released by the sensors at the bottom of the sensor plate wells and converting it to accurate OCR data points. Most currently available systems lack the capability to include kLa into the processing step, severely limiting their culturing duration and flexibility due to lack of continuous mixing function and intra-well gradients. Its inclusion was particularly important to the DolphinQ system, as users can change between different plate and mixing formats, affecting oxygen uptake and natural acidification rate. To reduce natural variance and noise sensitivity, multiple pre-processing steps are incorporated into the OCR and ECAR computation. Most notably, locally weighted scatterplot smoothing, better known as LOWESS, was applied at 0.2 strength, with higher strength values indicating increased strictness of the smoothening (Figure 3). Other data adjustments, such as weighted averages and baseline corrected readouts can be chosen by the user through the device interface. The baseline correction function can be adjusted mid culture to observe how growth and metabolic performance correlates to different controls or blanks, reducing the need for additional imaging and providing easily understandable visualizations of the experiment trajectory. The raw data obtained can further be used to calculate metabolic performance for only specific time-points during the culture or to understand OCR/ECAR per cell [^32^].

**Table 1.**
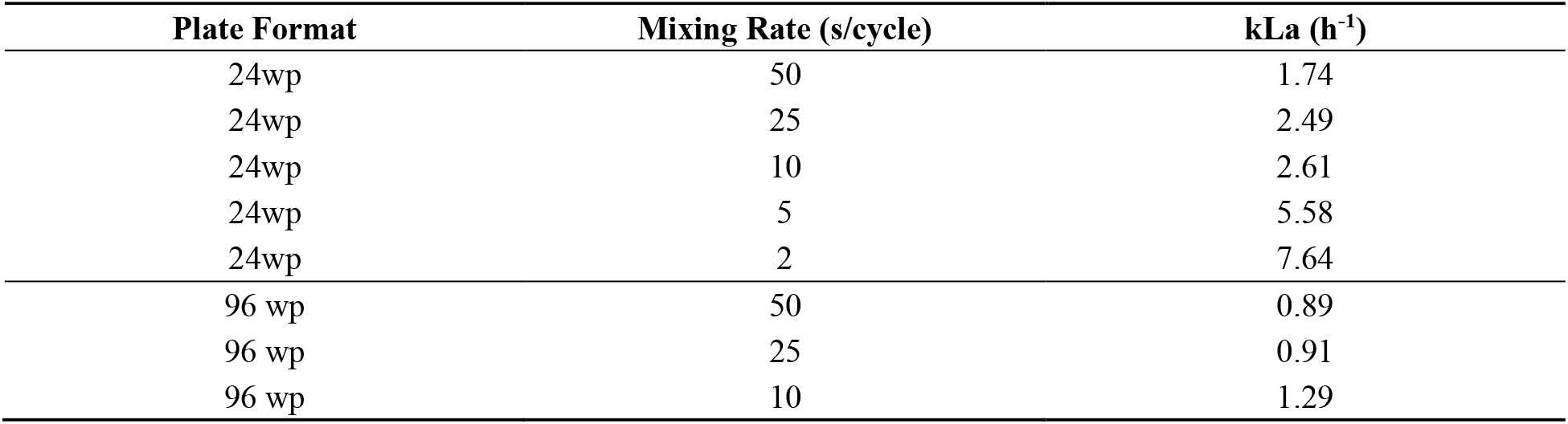
The measured oxygen transfer coefficients (kLa) under different plate formats and mixing rates.

| Plate Format | Mixing Rate (s/cycle) | kLa (h<sup>-1</sup>) |
| --- | --- | --- |
| 24wp | 50 | 1.74 |
| 24wp | 25 | 2.49 |
| 24wp | 10 | 2.61 |
| 24wp | 5 | 5.58 |
| 24wp | 2 | 7.64 |
| 96 wp | 50 | 0.89 |
| 96 wp | 25 | 0.91 |
| 96 wp | 10 | 1.29 |

The project was further supported with readouts obtained on several commonly used cell types and a case study using cells with mtDNA mutations affecting the metabolic performance of affected cells. Results showed clear, seeding density dependent performance and low inter-groups variance especially on lower seeded groups (Figure 4). An additional experimental design was developed, involving cells harboring a common Leigh syndrome mtDNA mutation. The disease model was chosen as it had clearly defined subgroups with well characterized metabolic profiles. Mitochondrial diseases are amongst the most complex, as they influence all aspects of cellular growth and performance, making them harder to dissect with conventional technologies. As predicted, the Rho0 cells and mutant type cells showed insufficiencies in OXPHOS and a normal or improved glycolytic profile observable in both pH and ECAR. These findings were replicated in the 96-well format providing valuable insight into how total cell number and plate format notably change the raw consumption numbers, yet not the trends and accuracy of the readout. Furthermore, a primary human fibroblast model also harboring the mtDNA mutation was shown to be visibly inhibited by oligomycin, a potent ATPase targeting toxin (Figure 6). Every cell in our body has slightly different metabolic tendencies and the gaps widen even further when immortalization or cancer derived cell lines are involved [^33,34,35^]. To understand these changes, prolonged observation of growth and maturation in cell types is essential and has been a bottleneck in the field.

This study supports the use of the DolphinQ system in long-term metabolic profiling and general cell culture. Future experiments will involve additional primary and non-mammalian cell lines, as well as more model systems such as spheroids and organoids.

## 5. Conclusions

The DolphinQ bioanalyzer bridges the gap between high-throughput metabolic flux analysis and long-term cell culture. By accounting for the physical dynamics of oxygen transfer through agitation and incorporating sophisticated data-smoothing algorithms, the system provides a high-fidelity narrative of cellular bioenergetics. By accounting for the physical dynamics of oxygen transfer through agitation and incorporating sophisticated data-smoothing algorithms, the system provides a high-fidelity narrative of cellular bioenergetics. This technology is particularly suited for studies requiring the long-term observation of metabolic transitions, such as disease modeling, stem cell differentiation, and pharmacological screens.

## Author Contributions

Conceptualization, D.L. and D.H.K.; methodology, D.L. Q.L.L. and D.H.K.; validation, D.H.K., Q.L.L.; investigation, D.L., S.H. and D.H.K.; data curation, D.L., Q.L.L.., D.H.K .; writing—original draft preparation, D.L. and D.H.K.; writing—review and editing, D.L., Q.L.L and D.H.K.; visualization, D.L. and D.H.K.; supervision, C.H.T., D.L. Q.L.L.and D.H.K.; project administration, D.L. and C.H.T.; All authors have read and agreed to the published version of the manuscript.

## Abbreviations

The following abbreviations are used in this manuscript:

A375: Human malignant melanoma cell line
A549: Human lung adenocarcinoma epithelial cell line
ATP: Adenosine triphosphate
C3A: Subclone of HepG2 human hepatocellular carcinoma cells
CO_2_: Carbon dioxide
ddH_2_O: Double-distilled water
ddPCR: Droplet digital Polymerase Chain Reaction
DMEM: Dulbecco’s Modified Eagle Medium
DNA: Deoxyribonucleic acid
DO: Dissolved oxygen in liquid media
DPBS: Dulbecco’s phosphate-buffered saline
ECAR: Extracellular acidification rate
FADH_2_: Flavin adenine dinucleotide (Fully reduced, dihydrogenated form)
HepG2: Human hepatocellular carcinoma cell line
kLa: Volumetric mass transfer coefficient for oxygen
LHON: Leber hereditary optic neuropathy
LOWESS: Locally Weighted Scatterplot Smoothing
mtDNA: Mitochondrial Deoxyribonucleic Acid
mt-ISR: Mitochondrial integrated stress response
MT-ATP6: Mitochondrially Encoded ATP Synthase Membrane Subunit 6
nDNA: Nuclear Deoxyribonucleic Acid
NADH: Nicotinamide adenine dinucleotide (Reduced form)
NaHCO_3_: Sodium bicarbonate
NEAA: Non-essential amino acids
OCR: Oxygen consumption rate
OTR: Oxygen transfer rate
OXPHOS: Oxidative phosphorylation
pH: Potential of hydrogen
SEM: Standard error of the mean
UV: Ultraviolet radiation
VAF: Variant allele frequency

## References

(1) Torralba, D.; Baixauli, F.; Sanchez-Madrid, F. Mitochondria Know No Boundaries: Mechanisms and Functions of Intercellular Mitochondrial Transfer. Front Cell Dev Biol 2016, 4, 107. DOI: 10.3389/fcell.2016.00107

(2) Roger, A. J.; Munoz-Gomez, S. A.; Kamikawa, R. The Origin and Diversification of Mitochondria. Curr Biol 2017, 27 (21), R1177–R1192. DOI: 10.1016/j.cub.2017.09.015

(3) Liberti, M. V.; Locasale, J. W. Metabolism: A new layer of glycolysis. Nat Chem Biol 2016, 12 (8), 577–578. DOI: 10.1038/nchembio.2133

(4) Spinelli, J. B.; Haigis, M. C. The multifaceted contributions of mitochondria to cellular metabolism. Nat Cell Biol 2018, 20 (7), 745–754. DOI: 10.1038/s41556-018-0124-1

(5) Smith, R. L.; Soeters, M. R.; Wust, R. C. I.; Houtkooper, R. H. Metabolic Flexibility as an Adaptation to Energy Resources and Requirements in Health and Disease. Endocr Rev 2018, 39 (4), 489–517. DOI: 10.1210/er.2017-00211

(6) Tsilingiris, D.; Tzeravini, E.; Koliaki, C.; Dalamaga, M.; Kokkinos, A. The Role of Mitochondrial Adaptation and Metabolic Flexibility in the Pathophysiology of Obesity and Insulin Resistance: an Updated Overview. Curr Obes Rep 2021, 10 (3), 191–213. DOI: 10.1007/s13679-021-00434-0

(7) Olson, K. A.; Schell, J. C.; Rutter, J. Pyruvate and Metabolic Flexibility: Illuminating a Path Toward Selective Cancer Therapies. Trends Biochem Sci 2016, 41 (3), 219–230. DOI: 10.1016/j.tibs.2016.01.002

(8) Thorburn, D. R. Mitochondrial disorders: prevalence, myths and advances. J Inherit Metab Dis 2004, 27 (3), 349–362. DOI: 10.1023/B:BOLI.0000031098.41409.55

(9) Gorman, G. S.; Chinnery, P. F.; DiMauro, S.; Hirano, M.; Koga, Y.; McFarland, R.; Suomalainen, A.; Thorburn, D. R.; Zeviani, M.; Turnbull, D. M. Mitochondrial diseases. Nat Rev Dis Primers 2016, 2, 16080. DOI: 10.1038/nrdp.2016.80

(10) Wen, H.; Deng, H.; Li, B.; Chen, J.; Zhu, J.; Zhang, X.; Yoshida, S.; Zhou, Y. Mitochondrial diseases: from molecular mechanisms to therapeutic advances. Signal Transduct Target Ther 2025, 10 (1), 9. DOI: 10.1038/s41392-024-02044-3

(11) Huss, J. M.; Kelly, D. P. Mitochondrial energy metabolism in heart failure: a question of balance. J Clin Invest 2005, 115 (3), 547–555. DOI: 10.1172/JCI24405

(12) Wang, Z.; Ying, Z.; Bosy-Westphal, A.; Zhang, J.; Schautz, B.; Later, W.; Heymsfield, S. B.; Muller, M. J. Specific metabolic rates of major organs and tissues across adulthood: evaluation by mechanistic model of resting energy expenditure. Am J Clin Nutr 2010, 92 (6), 1369–1377. DOI: 10.3945/ajcn.2010.29885

(13) Gorlach, A.; Bertram, K.; Hudecova, S.; Krizanova, O. Calcium and ROS: A mutual interplay. Redox Biol 2015, 6, 260–271. DOI: 10.1016/j.redox.2015.08.010

(14) De Nicolo, B.; Cataldi-Stagetti, E.; Diquigiovanni, C.; Bonora, E. Calcium and Reactive Oxygen Species Signaling Interplays in Cardiac Physiology and Pathologies. Antioxidants (Basel) 2023, 12 (2). DOI: 10.3390/antiox12020353

(15) Green, D. R.; Kroemer, G. The pathophysiology of mitochondrial cell death. Science 2004, 305 (5684), 626–629. DOI: 10.1126/science.1099320

(16) Pakos-Zebrucka, K.; Koryga, I.; Mnich, K.; Ljujic, M.; Samali, A.; Gorman, A. M. The integrated stress response. EMBO Rep 2016, 17 (10), 1374–1395. DOI: 10.15252/embr.201642195

(17) Fu, Y.; Sacco, O.; DeBitetto, E.; Kanshin, E.; Ueberheide, B.; Sfeir, A. Mitochondrial DNA breaks activate an integrated stress response to reestablish homeostasis. Mol Cell 2023, 83 (20), 3740–3753 e3749. DOI: 10.1016/j.molcel.2023.09.026

(18) Liu, H.; Wang, S.; Wang, J.; Guo, X.; Song, Y.; Fu, K.; Gao, Z.; Liu, D.; He, W.; Yang, L. L. Energy metabolism in health and diseases. Signal Transduct Target Ther 2025, 10 (1), 69. DOI: 10.1038/s41392-025-02141-x

(19) Yoo, I.; Ahn, I.; Lee, J.; Lee, N. Extracellular flux assay (Seahorse assay): Diverse applications in metabolic research across biological disciplines. Mol Cells 2024, 47 (8), 100095. DOI: 10.1016/j.mocell.2024.100095

(20) Dranka, B. P.; Benavides, G. A.; Diers, A. R.; Giordano, S.; Zelickson, B. R.; Reily, C.; Zou, L.; Chatham, J. C.; Hill, B. G.; Zhang, J.; et al. Assessing bioenergetic function in response to oxidative stress by metabolic profiling. Free Radic Biol Med 2011, 51 (9), 1621–1635. DOI: 10.1016/j.freeradbiomed.2011.08.005

(21) Giglio, E.; Giuseffi, M.; Picerno, S.; Sichetti, M.; Mecca, M. Optimised Workflows for Profiling the Metabolic Fluxes in Suspension vs. Adherent Cancer Cells via Seahorse Technology. Int J Mol Sci 2024, 26 (1). DOI: 10.3390/ijms26010154

(22) Zhang, J.; Nuebel, E.; Daley, G. Q.; Koehler, C. M.; Teitell, M. A. Metabolic regulation in pluripotent stem cells during reprogramming and self-renewal. Cell Stem Cell 2012, 11 (5), 589–595. DOI: 10.1016/j.stem.2012.10.005

(23) Schmidt, C. A.; Fisher-Wellman, K. H.; Neufer, P. D. From OCR and ECAR to energy: Perspectives on the design and interpretation of bioenergetics studies. J Biol Chem 2021, 297 (4), 101140. DOI: 10.1016/j.jbc.2021.101140

(24) Bavli, D.; Prill, S.; Ezra, E.; Levy, G.; Cohen, M.; Vinken, M.; Vanfleteren, J.; Jaeger, M.; Nahmias, Y. Real-time monitoring of metabolic function in liver-on-chip microdevices tracks the dynamics of mitochondrial dysfunction. Proc Natl Acad Sci U S A 2016, 113 (16), E2231–2240. DOI: 10.1073/pnas.1522556113

(25) Narayanan, H.; Hinckley, J. A.; Barry, R.; Dang, B.; Wolffe, L. A.; Atari, A.; Tseng, Y. Y.; Love, J. C. Accelerating cell culture media development using Bayesian optimization-based iterative experimental design. Nat Commun 2025, 16 (1), 6055. DOI: 10.1038/s41467-025-61113-5

(26) Shelton, S. D.; House, S.; Martins Nascentes Melo, L.; Ramesh, V.; Chen, Z.; Wei, T.; Wang, X.; Llamas, C. B.; Venigalla, S. S. K.; Menezes, C. J.; et al. Pathogenic mitochondrial DNA mutations inhibit melanoma metastasis. Sci Adv 2024, 10 (44), eadk8801. DOI: 10.1126/sciadv.adk8801

(27) Shin, W. S.; Lee, D.; Kim, S.; Jeong, Y. S.; Chun, G. T. Application of scale-up criterion of constant oxygen mass transfer coefficient (kLa) for production of itaconic acid in a 50 L pilot-scale fermentor by fungal cells of Aspergillus terreus. J Microbiol Biotechnol 2013, 23 (10), 1445–1453. DOI: 10.4014/jmb.1307.07084

(28) Cleveland, W. S. Robust Locally Weighted Regression and Smoothing Scatterplots. Journal of the American Statistical Association 1979, 74 (368). DOI: 10.2307/2286407.

(29) Charis- P. Segeritz, L. V. Cell Culture: Growing Cells as Model Systems In Vitro; Academic Press, 2017. DOI: 10.1016/B978-0-12-803077-6.00009-6.

(30) Magliaro, C.; Mattei, G.; Iacoangeli, F.; Corti, A.; Piemonte, V.; Ahluwalia, A. Oxygen Consumption Characteristics in 3D Constructs Depend on Cell Density. Front Bioeng Biotechnol 2019, 7, 251. DOI: 10.3389/fbioe.2019.00251

(31) Garcia-Ochoa, F.; Gomez, E. Bioreactor scale-up and oxygen transfer rate in microbial processes: an overview. Biotechnol Adv 2009, 27 (2), 153–176. DOI: 10.1016/j.biotechadv.2008.10.006

(32) Kurucz, B.; Hajdinak, P.; Szarka, A. Resveratrol-Supported Bioenergetics Leads to Higher Productivity and Accompanying Endoplasmic Reticulum Stress in a mAb-Producing CHO Cell Line. Int J Mol Sci 2025, 26 (22). DOI: 10.3390/ijms262211146

(33) Savage, V. M.; Allen, A. P.; Brown, J. H.; Gillooly, J. F.; Herman, A. B.; Woodruff, W. H.; West, G. B. Scaling of number, size, and metabolic rate of cells with body size in mammals. Proc Natl Acad Sci U S A 2007, 104 (11), 4718–4723. DOI: 10.1073/pnas.0611235104

(34) Bordbar, A.; Feist, A. M.; Usaite-Black, R.; Woodcock, J.; Palsson, B. O.; Famili, I. A multi-tissue type genome-scale metabolic network for analysis of whole-body systems physiology. BMC Syst Biol 2011, 5, 180. DOI: 10.1186/1752-0509-5-180

(35) Gebert, N.; Rahman, S.; Lewis, C. A.; Ori, A.; Cheng, C. W. Identifying Cell-Type-Specific Metabolic Signatures Using Transcriptome and Proteome Analyses. Curr Protoc 2021, 1 (9), e245. DOI: 10.1002/cpz1.245

