## Supplemental Data 1-2 for "Long-term high throughput agitation culturing with real-time metabolic profiling"

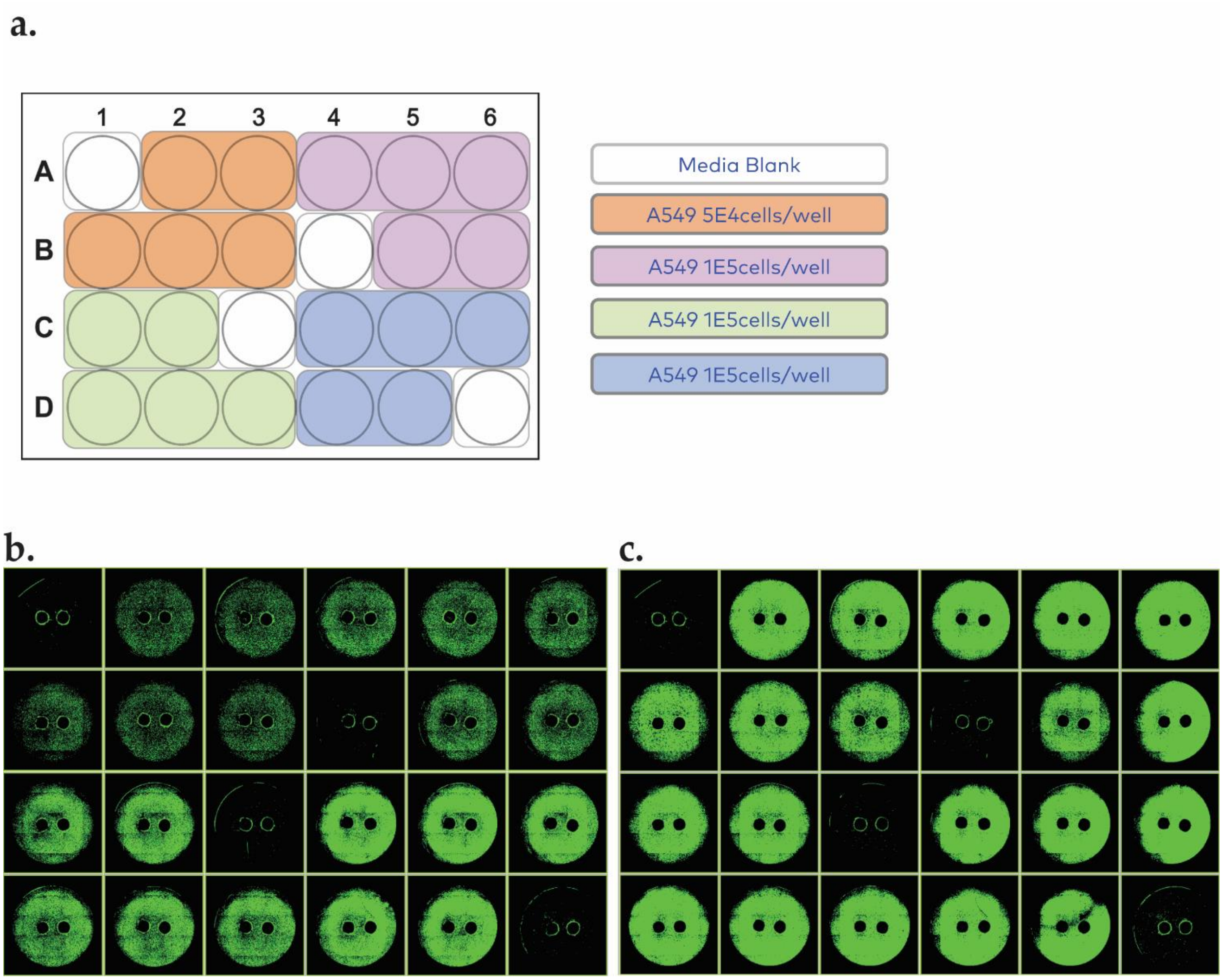


***Supplementary Figure 1.*** *Additional information on seeding and viable cell density (a) Plate layout of the Figure 4 experiment (b-c), Cell imager observations on Day 0 and Day 4 respecively*

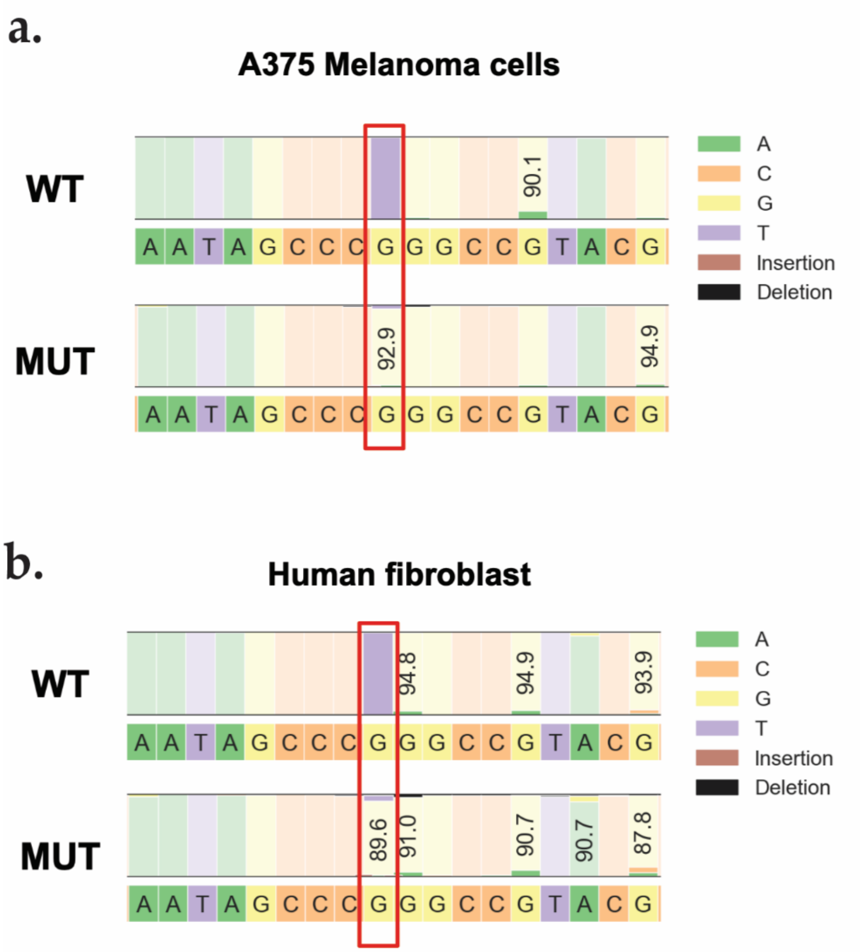


***Supplementary Figure 2.*** *MT-ATP6 heteroplasmy level in A375 cells (a) and human fibroblast cells (b). Red rectangle indicates m.8993 mutation site.*
